# The impact of phenological mismatch varies across woodland food-web interactions

**DOI:** 10.64898/2026.06.28.734640

**Authors:** Jamie C. Weir, Albert B. Phillimore

## Abstract

Climate warming is altering the timing of seasonal events across ecosystems, impacting the temporal synchrony of interactions among species^1,2^. For trophic interactions, the match-mismatch hypothesis predicts that when consumers become phenologically asynchronous with key ephemeral resources their fitness will decline^3–5^. Most studies of mismatch focus on single resource-consumer species pairs, and implicitly assume trophic specialisation. However, many consumers exploit more than one resource species, giving rise to several mechanisms whereby the negative impacts of mismatch on individuals and populations could be buffered^6^. Here we experimentally manipulate phenological asynchrony across 48 plant-caterpillar interactions in a spring woodland food-web system and assay caterpillar performance. As asynchrony increases, we find strong evidence for a decline in survival that generalises across host-caterpillar interactions, whereas caterpillar growth and development are largely unaffected. We also show that focus in the literature on a single model interaction (Oak-Winter Moth)^7,8^ has likely overestimated the general impact asynchrony in this system. The strength of the effect of mismatch varies markedly among host-plants, caterpillars, and their interactions—with a small number of interactions showing little or no decline in consumer performance despite substantial asynchrony. Our results demonstrate that the fitness consequences of phenological mismatch are widespread but interaction-specific, revealing substantial heterogeneity in how trophic interactions are expected to respond to climate-driven shifts in seasonal timing. This variation in response could allow resource diversity and resource switching to buffer consumer guilds against the phenological impacts of ongoing climate change, stabilising the abundance of caterpillars for higher trophic levels.

---

Phenological shifts are a conspicuous and well-documented effect of recent climate warming^9^. However, these shifts vary considerably in magnitude among species, guilds, and trophic levels^1,2^. For taxa that depend on phenological synchrony with other species, divergent responses could be particularly consequential for population size and even overall ecosystem resilience^10^. The match-mismatch hypothesis, first developed in the context of population recruitment in marine fishes^3,4,11^, is widely applied to systems where the degree of phenological synchrony between interacting species has fitness consequences for one or both parties. Negative impacts of phenological asynchrony on the survival and reproduction of mistimed species have been reported across a breadth of systems, including arboreal caterpillars that feed on the first flush of spring leaves^8^, passerine birds feeding on caterpillars^12^, cod feeding on zooplankton^5^, and flowering plants relying on bee pollinators^13^.

A central tenet of the match-mismatch hypothesis as applied to trophic systems is that the fitness of a specialist consumer is sensitive to the level of synchrony (co-incident timing) with a time-limited resource^14^. This theoretical context likely explains the tendency of field and laboratory studies of trophic mismatch to focus on simple food-chains^7,12,15^, often implicitly framing consumer species as specialists. In reality, many consumers actually feed on a number of different resource species, yet we are unaware of any studies examining the fitness consequences of phenological asynchrony within a multi-resource and multi-consumer framing. It is therefore unclear whether the negative fitness effects of asynchrony that have been reported for some simplified food-chains are representative, and whether they generalise across consumer species, resource species, or pairwise species combinations^6^. Where responses vary among species interactions, this has the potential to stabilise the impacts of environmental change that accrue at higher levels of organisation (e.g., population or food-web) through portfolio effects^16^ and/or compensatory dynamics^17,18^. Despite many case studies documenting the negative impacts of asynchrony on fitness, this is yet to translate into widespread evidence for population-level declines^14^. The response diversity of asynchrony-performance relationships, and emergent portfolio effects, are one potential explanation for this disconnect^6^.

## Mismatch in temperate woodland food-webs

The temperate woodland *tree*→*caterpillar*→*bird* food-chain has become a classic model system for studying phenological mismatch in the context of climate change^19,20^. The flush of new tree foliage in spring provides ‘matched’ (synchronous) phytophagous caterpillars with an abundant food resource. Caterpillars hatching or breaking winter diapause too early can starve^7,21,22^, and those hatching too late are forced to feed on older and better-defended foliage^23,24^. Caterpillar timing in spring is therefore thought to be under strong selection to match a narrow window of optimal host-plant availability^7,21^. The phenology of both leaf-flushing and caterpillar hatching/activity is sensitive to spring temperatures^20,25,26^, but differences in the phenological sensitivity to temperature of these two trophic levels has the potential to generate year-to-year variation in temporal synchronisation^26^, and hence fluctuations in the growth and survival of caterpillars^21^.

Caterpillars are abundant defoliators in spring woodlands and play a major role in nutrient cycling^27^. But while the arboreal caterpillar guild are species-rich^28^ and generally highly polyphagous^29,30^, previous work has focused almost exclusively on a single model trophic interaction—Winter Moth (*Operophtera brumata*) caterpillars feeding on young English Oak (*Quercus robur*) foliage. The Winter Moth has been recorded feeding on plants from 31 genera across 15 families, on species ranging from birch to spruce, heather, and bilberry ^31^. Asynchrony with Oak bud-burst has been widely reported to negatively impact Winter Moth caterpillar performance^7,21^, though the extent to which this single trophic interaction is representative of the diversity of interactions occurring across the wider food-web is unclear (i.e., alternative host-plants or other caterpillar species). Alongside **trophic generalism in caterpillars** and **variation in tree leafing phenology**, differences in baseline performance (i.e., where asynchrony = 0 days) among caterpillars, hosts, and pairwise interactions, and differences in the impacts of asynchrony on fitness (i.e., the slope of performance vs hatching time), could buffer individuals and populations through a combination of **host switching** (where young caterpillars disperse to alternate hosts) and **portfolio effects** (that optimise geometric mean performance at the aggregate level) (Fig 1a)^6^. Collectively, by buffering total caterpillar abundance against asynchrony, these mechanisms could also stabilise top-down effects on host-plants and bottom-up effects on secondary consumers, such as birds.

**Fig 1.**
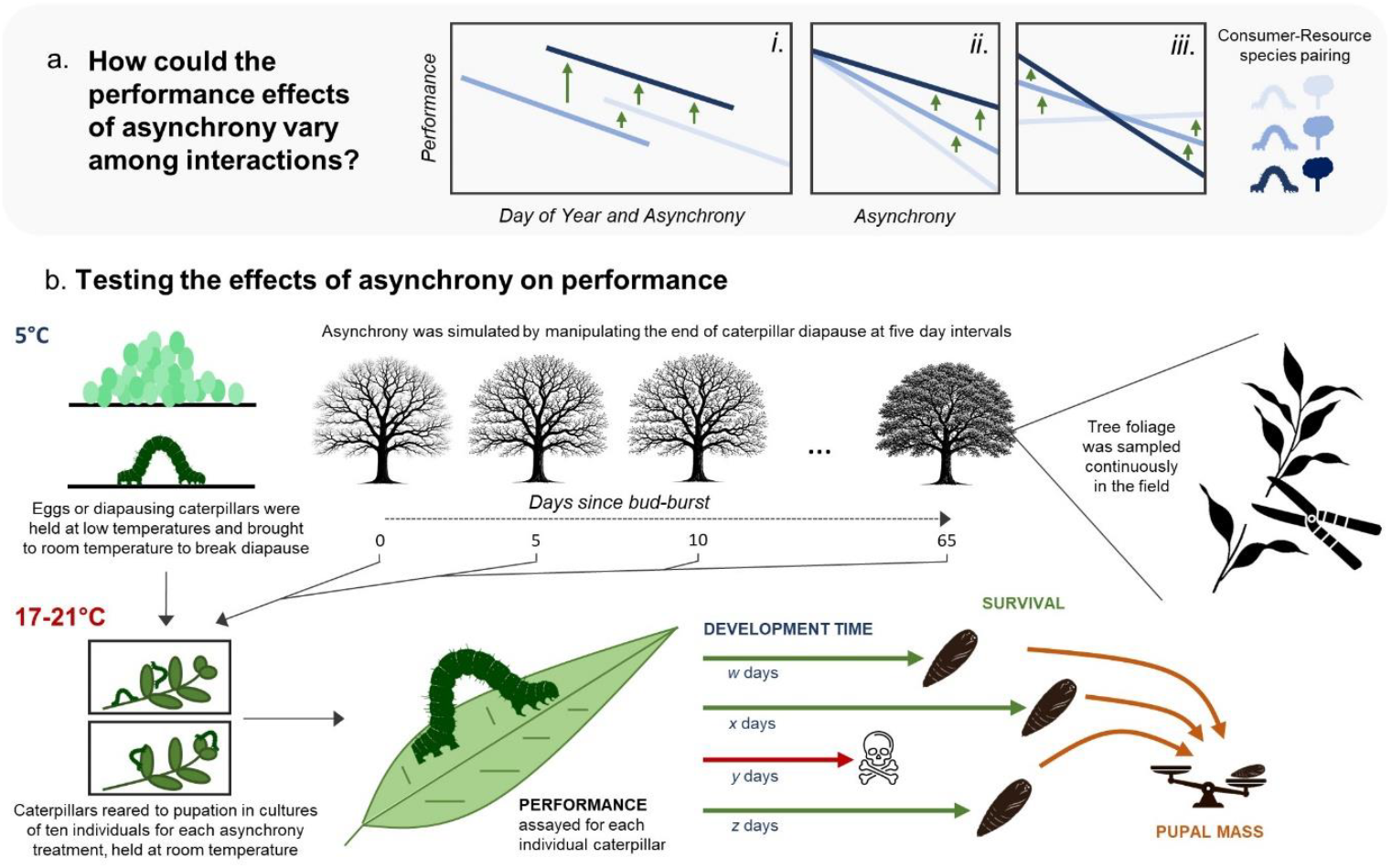
**(a) Schematic showing three hypothetical patterns of among resource-consumer variation in the effects of asynchrony on caterpillar (consumer) performance**: (*i*) same relationship (slope) over time, but variation in baseline performance on that host (intercept); (*ii*) variation in the relationship over time, but common baseline performance; (*iii*) variation in both the relationship and baseline. A generalist diet can facilitate buffering in each case, via host switching at the individual level and/or portfolio effects at the population level. For any given value of asynchrony, generalism gives an individual caterpillar the possibility of increasing its performance by utilising an alternative resource. At the aggregate (population) level, where individual caterpillars are spread across different hosts, variation (both in the baseline and the slope of the asynchrony relationship) will minimise year-to-year variation in average performance. Even where the slope of the performance-asynchrony relationship is equivalent (plot *i*), variation in baseline performance on a given host coupled with variation in the spring phenology of different host and caterpillar species (the ‘Day of Year’ start-date of each line in *i* means that the optimal host species can still vary temporally. **(b) Experimental overview**: to explore variation in the performance effects of asynchrony across trophic interactions, we reared six caterpillar (consumer) species (Winter Moth *Operophtera brumata*, Mottled Umber *Erannis defoliaria*, Gypsy Moth *Lymantria dispar*, Black Arches *Lymantria monacha*, the Vapourer *Orgyia antiqua*, and Scarce Vapourer *Orgyia recens*) on eight host-plant (resource) species (Alder *Alnus glutinosa*, Apple *Malus domestica*, Birch *Betula pendula*, Hawthorn *Crataegus monogyna*, Oak *Quercus robur*, Sallow *Salix capraea*, Sycamore *Acer pseudoplatanus*, and Willow *Salix alba*). For each host-caterpillar species pair, asynchrony was simulated at intervals of five calendar days (from 0-65 days) by manipulating the timing of caterpillar diapause break. Caterpillars were reared in ‘cultures’ of ten individuals and we assayed three metrics of individual performance: survival to pupation, final pupal mass, and development time from egg hatch to pupation.

Here, we present the first experimental study assaying the performance impacts of trophic asynchrony replicated across a web of resource-consumer interactions. We reared over 20,000 caterpillars of six spring arboreal species on eight of their common native host-plant species (Fig 1b, S1). We examined how three components of individual performance—survival, growth, and time to pupation—were impacted by asynchrony with leaf bud-burst. Asynchrony was manipulated by hatching (or breaking diapause for) caterpillars on different dates relative to their resource taxon (Fig 1b). We aimed to: **(1)** Establish whether a negative impact of trophic asynchrony on consumer performance is a general property across resource and consumer taxa in this food-web; **(2)** Examine the extent to which performance at synchrony and with increasing asynchrony varies among host-plant species, caterpillar species, and their unique combination; **(3)** Consider the implications of asynchrony response diversity for mechanisms whereby host-plant generalism may buffer the impacts of phenological asynchrony on caterpillar performance, and also stabilise the effect of host-by-caterpillar asynchrony on higher and lower trophic levels.

### Mismatch is general, but the effects are interaction specific

To examine the effects of asynchrony on performance we used linear mixed models (LMMs) and generalised mixed models (GLMMs) that include all resource-consumer species pairs, with host-plant, caterpillar species and their interaction included as random intercept and slope terms. Across the 48, host-by-caterpillar combinations, the average trend was for the proportion of caterpillars surviving to pupation to decline with increasing asynchrony (binomial response GLMM slope = −0.05 log odds/day, 95% credible intervals [hereafter CI]: −0.08 — −0.03), from a proportion of 0.58 (0.40 — 0.75) at complete synchrony to 0.24 (CI: 0.09 — 0.42) with 65 days of asynchrony (Fig 2). Contrary to expectations, neither pupal mass (Gaussian LMM slope = −0.0006 mg/day, CI: −0.0085 — 0.0078, Fig 2) nor time to pupation (Gaussian LMM slope = 0.03 days/day, CI: −0.15 — 0.21, Fig 2) decreased significantly as asynchrony increased (Tables S1-S3) Given that survival appears to be the performance metric that is most sensitive to asynchrony, we focus most of our attention on this trait. Surprisingly, we find that the average impact of asynchrony on survival was quite shallow, extending over a much more protracted timeframe than has been reported in Oak-Winter Moth studies ^7,21^—on a scale of weeks, rather than days.

**Fig 2.**
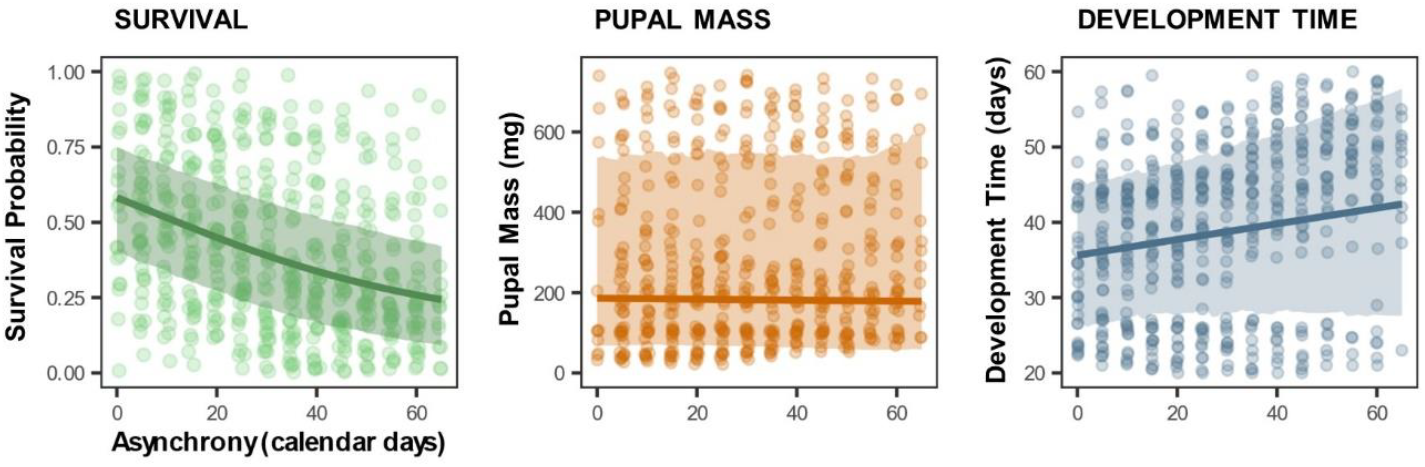
The predicted effects of asynchrony on caterpillar survival to pupation, pupal mass, and development time to pupation. Solid lines are the posterior mean of the average effect across all host-caterpillar interactions, with shaded 95% credible intervals. Points show averages of raw data for each host-caterpillar combination in each asynchrony treatment.

For all three performance metrics, we found that most of the variance (for survival this was assessed on the latent scale) was due to variation in the intercept of the performance-asynchrony relationship (survival = 70%, CI: 47— 90; mass = 78%, CI: 54 — 96; duration = 66%, CI: 35 — 91; Fig 3; Table S4). For survival, a substantial proportion of the intercept variance was accounted for by caterpillar species (22%, CI: 0 — 61), host-plant species (13%, CI: 0 — 44), and their interaction (34%, CI: 11 — 60), though credible intervals for caterpillar and host-plant variance are not distinguishable from 0. These variance estimates reveal that when hatching is synchronous with the timing of bud-burst, some caterpillar species may survive better than others, survival may be higher on some host-plants than others, and certain caterpillar species clearly survive particularly well (or poorly) on certain host-plants. For both mass (58%, CI: 0 — 90) and larval duration (43%, CI: 0 — 80), most of the intercept variance is attributable to among-caterpillar species differences, though the credible intervals are very broad. A significant proportion of the variance in survival arises from variation in the effect of asynchrony (16%, CI: 4 — 38)—i.e., variance in slope. Much of the asynchrony-survival slope variance is attributable to differences between caterpillar species (46%, CI: 9 — 87) and host-plants (38%, CI: 7 — 74), with the variance in the caterpillar-by-host interaction smaller and not distinguishable from 0 (6%, CI: 0 — 34).

**Fig 3.**
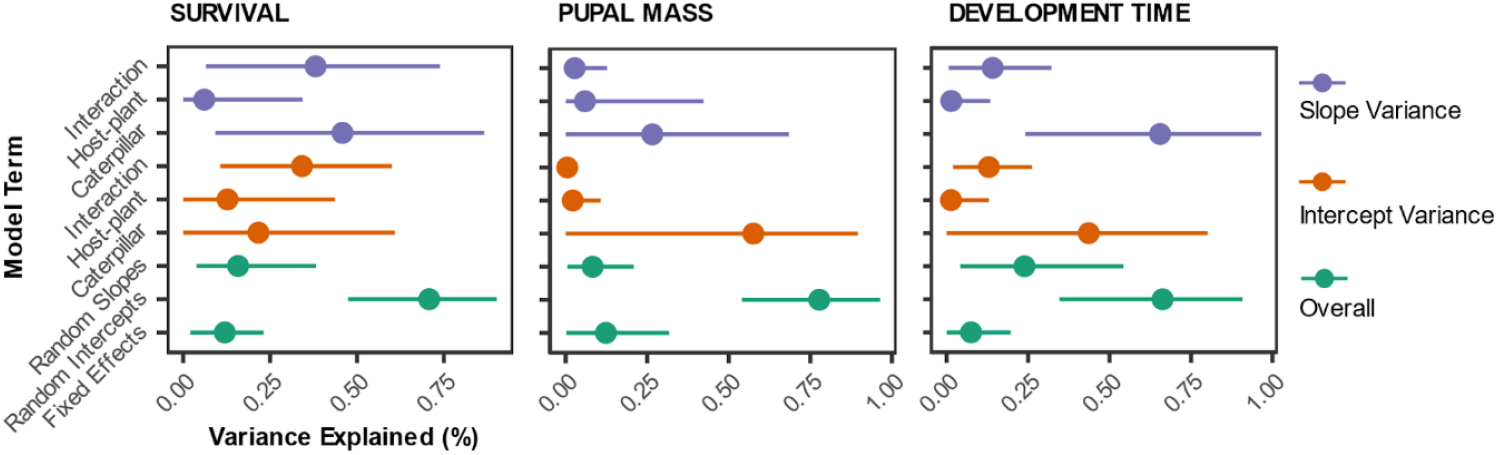
Variance components of three metrics of caterpillar performance with increasing asynchrony. Overall variance is partitioned to show relative contribution of fixed effects, random slopes, and random intercepts 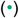. Variance is partitioned to show the relative contribution of among-caterpillar, among-host-plant, and the host-plant-by-caterpillar interaction to the random slopes 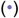 and random intercepts 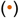. The level at which most variation exists in the effects of asynchrony affects our interpretation of the mechanisms that can contribute to buffering in this system (see Figure 1a).

Across caterpillar species we observe a general tendency for some species to survive better than others on the average host-plant, with the Vapourer exhibiting highest survival and Winter Moth the lowest (Fig 4a). A negative effect of asynchrony on caterpillar survival was significant for all six caterpillar species (Table S5), with the decline in survival as asynchrony increases being steepest for Winter Moth (Fig 4a; slope = −0.08 log odds/day, CI: −0.10 — −0.06) and departing significantly from the general trend (average slope = −0.05 log odds/day, CI: −0.07— −0.03). This suggests that a disproportionate focus on using the Winter Moth as a representative model species in mismatch research^7,15,21,32,33^ may overstate the wider performance impacts of asynchrony across the guild.

**Fig 4.**
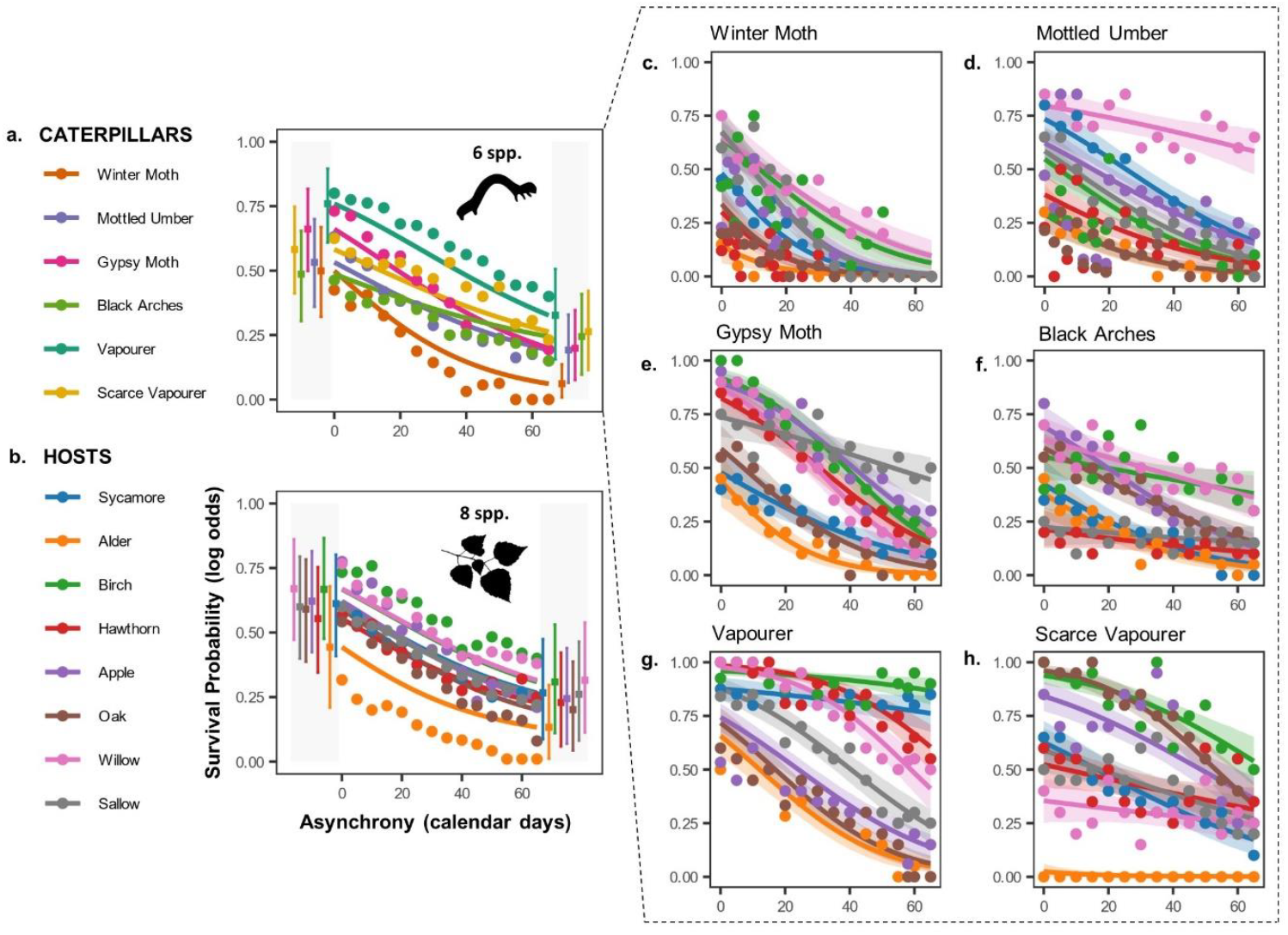
Average predicted effects of asynchrony on survival to pupation across (a) six spring caterpillar species and (b) eight of their host-plant species. Inset panel (c-h) shows the predicted relationships in all pairwise combinations of each caterpillar species fed on each host-plant. Trends represent posterior mean model estimates with shaded 95% credible intervals. Square points in the grey shaded zone of the species average plots (a and b) correspond to model estimates (± 95% CIs) of survival probability at the minimum (0 days) and maximum (65 days) values of induced asynchrony. Round points on all panels show numerical averages of raw data for each level of asynchrony across the relevant groups.

We find that the significant negative effect of asynchrony on the survival of the average caterpillar species is replicated across all eight tree species (Fig 4b; Table S5). The most striking difference among tree species was that survival was consistently higher on some, such as Birch and Willow, and consistently low on Alder^33^, which partly agrees with differences in the observed abundance of caterpillars on different hosts in the field^34^. Among-host differences in the effects of asynchrony on survival of the average caterpillar species are small, such that the same host species is generally best (or worst) regardless of the degree of asynchrony. For Oak, frequently a focal species in mismatch studies^7,21,22,24,26^, we find that survival of the average caterpillar (0.59, CIs: 0.39 — 0.78) is close to the overall average at an asynchrony of 0 (0.47, CIs: 0.29 — 0.64), but that the decline in survival on this host-plant is steeper than for the average tree taxon (slope = −0.06 log odds/day, CI: −0.09 — −0.03), consistent with the leaves of this species acquiring defences as they age^23^.

For 44 out of 48 host-by-caterpillar interactions, we identified a significant negative effect of asynchrony on survival (Fig 4c-h, 5; Table S6). The exceptions are the Black Arches on three of its host-plants (Hawthorn, Birch, and Sallow) and the Scarce Vapourer on White Willow—in these cases, we found no effect of asynchronous hatching on caterpillar survival. When reared on Alder, the Scarce Vapourer suffered complete mortality at all levels of asynchrony, and we conclude they cannot subsist on this host-plant. For the Oak-Winter Moth interaction, so prominent in mismatch research, our point estimate for the asynchrony slope is more steeply negative than all but one of the remaining 47 interactions (Fig 5).

**Fig 5.**
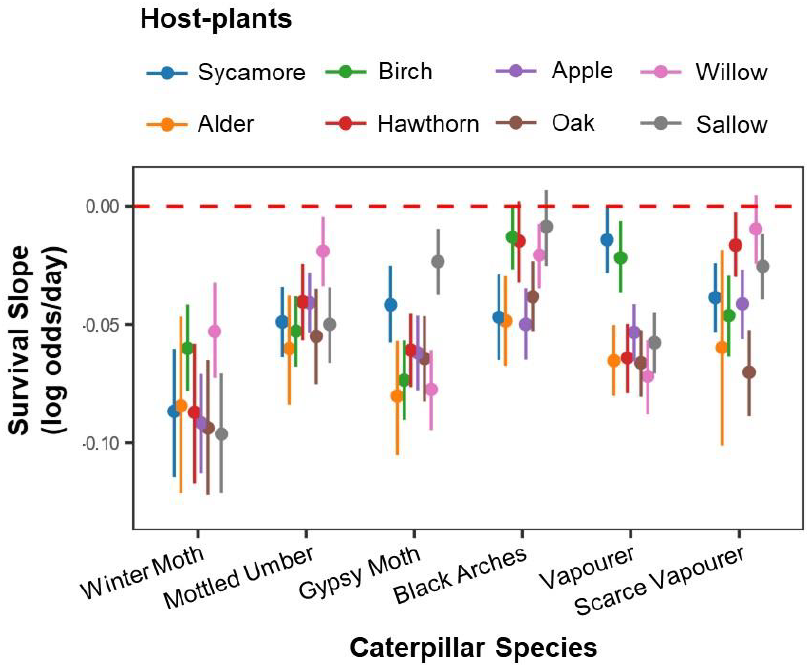
Slope estimates for the effect of asynchrony on caterpillar survival probability from egg hatch to pupation across each host-caterpillar interaction. Points correspond to posterior means with 95% credible interval bars.

Across the experiment, we can identify two main mechanisms whereby trophic generalism can buffer the impacts of phenological mismatch on caterpillars. **First**, we find some evidence that the host-plant that maximises performance changes as asynchrony increases, providing conditions that would allow for buffering via host-switching or host generalism—for example, survival of the average caterpillar is higher on Apple than Sallow when hatching is synchronous (Fig 4b), but this is reversed as asynchrony increases. We find a similar opportunity for host-switching to buffer mismatch that is particular to specific caterpillar species. For instance, at asynchrony of 0 days, Gypsy Moths survive best on Birch (0.92, CI: 0.87 — 0.96), whereas at asynchrony of 65 days survival is highest on Sallow (0.44, 0.34 — 0.55), as survival declines less steeply on this host plant (Fig. 3e). For Black Arches, in contrast, survival is highest on Apple at asynchrony of 0 days (0.69, CIs: 0.59 — 0.79), declining to (0.13, CIs: 0.07, 0.19) by 65 days—for individuals feeding on Birch, survival is substantially higher at 65 days (0.38, CIs: 0.28 — 0.49) (Fig 4f). **Second**, buffering can arise through inherent variation in host-plant leaf-out phenology (Roberts et al. 2016). This variation means that, in any given year, at least some caterpillars are likely to be synchronous with some host-plants, generating a portfolio effect at the population level that reduces the impact of asynchrony on interannual variation in population size.

Notably, the diversity of responses to asynchrony that we observe across host-plant-caterpillar interactions itself has the potential to serve as a buffer on the impacts of mismatch at higher trophic levels, by conferring greater stability on the availability of caterpillars to generalist secondary consumers and parasitoids (Fig 4c-h). Similarly, this response diversity is expected to stabilise the impact of herbivory on trees.

### Effects of asynchrony on pupal mass and caterpillar development time

Prior work on the Winter Moth has suggested that, whereas early hatching caterpillars lack food and starve^22^, the main contributor to declining fitness associated with late hatching asynchrony is lower leaf nutritional quality resulting in smaller pupae^7,35^, and subsequently less fecund adult females^33^. However, here we find that survival is the fitness component most sensitive to asynchrony, across host and caterpillar species. Although there is no significant overall trend in the effect of asynchrony on pupal mass (Table S2, S7), some host-by-caterpillar combinations do show trends that depart from a slope of zero. Where significant patterns exist, these are predominantly negative (5 of 48 caterpillar-by-host relationships), with only a single positive slope across all combinations (Fig S2; Table S8).

The duration of time spent in the larval stage is also expected to impact fitness, though reasonable predictions can be made in either direction: a shortened development time could be either a sign of stress in caterpillars and a strategy to avoid harsh/unfavourable conditions, or a sign of favourable conditions that promote rapid growth^e.g.36–41^. Temporal exposure to predation, parasitism, and disease across the larval and pupal periods will dictate the stage in which it is best to spend more life history time—a balance that is very likely to vary among taxa. As with pupal mass, although there is no significant overall effect of asynchrony on development (Table S3), the effect of asynchrony is significant for 29 out of 96 host-by-caterpillar-by-sex relationships, and in each case the duration of the larval period increases with asynchrony (Table S9-S10). Notably, in the Scarce Vapourer, increased asynchrony lengthened development time in both males and females, across all host-plant species (Fig S3).

## Discussion

Here we present the first experimental examination of the ecological importance of mismatch and its ubiquity across a diverse consumer guild. We find that there is a general impact of asynchrony on performance—particularly survival—across the interactions we considered, consistent with this being a general feature of interactions in this temperate forest food-web. While insights into mismatch from the Oak-Winter Moth interaction are pervasive in the literature^7,21^, we find that the steepness with which survival declines in this interaction is unrepresentative of the shallower declines found on average across the other host-caterpillar interactions. Overall, the impact of asynchrony on performance observed in this study is substantially less severe than has been reported elsewhere^7^, and is mediated via impacts on survival, rather than growth or development time. Taken together, the rather shallow impact of asynchrony on performance and the potential for caterpillar species to buffer asynchrony or its impacts by utilising different host-plants may contribute to many such species being more resilient to future climate change than has been appreciated hitherto^6^.

Our experiments reveal that the impacts of asynchrony on performance cannot be predicted solely on the basis of the average response of particular consumer and resource species, but rather that a substantial part of those impacts are interaction-specific. We might observe this where, for example, different consumers are adapted to different hosts^29^, and differ in their sensitivity to various host defences^42,43^. Within woodland ecosystems, the variation we observe in the impacts of asynchrony across interactions and the buffering effects this can generate are important considerations both for assessing the resilience of natural forest communities under climate change, and for engineering resilience in man-made natural or economically important forests. More generally, widespread buffering of the kind we explore in this study has the potential to resolve a persistent paradox in global change ecology: that while the negative effects of asynchrony are often evident for consumers at the level of the individual, they are much less clear, and certainly less documented, at the level of consumer populations^14,44^. It remains to be established, however, whether the broad insights gained here regarding the generality and/or specificity of the impacts of asynchrony generalise to other food-webs, ecosystems, or interaction network types.

Within the wider context of phenological mismatch, experimental manipulations of asynchrony remain unusual (but see^21,45^. Most of our insights into trophic mismatch still come from observational studies that are subject to confounding variables—for example, the potential confounding effect of individual quality on timing and fitness^46^. While for many systems a clean manipulation of timing is notoriously challenging^46^, the host-caterpillar interaction presents a particularly promising arena for experimental manipulations of asynchrony, by manipulating the timing of caterpillar diapause break^7,21^. Here we have demonstrated that a manipulation that has often been confined to the Winter Moth on Oak can be replicated across a wide range of caterpillar and plant species.

To limit the complexity of our experimental design we held temperature constant, though temperature is expected to have a direct impact on measures of performance^47^. We cannot exclude the possibility that the impacts of asynchrony interact with temperature, and that the interaction effect could potentially vary across host-caterpillar trophic interactions. As a result, given that our experiment temperature of ~19°C is higher than ambient spring temperature in the UK, some caution is required if extrapolating our laboratory findings to specific interactions in a natural context. Our use of constant temperature also departs from the natural fluctuations in day/night temperatures that are likely to affect growth rates^48^ and there would be value in repeating these experiments at a range of natural temperatures to pick apart the relative contribution of temperature and asynchrony under climate change. It is also possible that absolute timing has a negative impact on consumer performance, such that the caterpillars for which we induced a delay in hatching or breaking diapause would suffer a reduction in performance. However, this cannot account for the host-caterpillar combinations where we observe no significant decline in performance with asynchrony. Therefore, if such an effect exists, it cannot be universal across caterpillar taxa.

The diversity of responses to asynchrony we find across caterpillars, host-plants, and pairwise interactions demonstrate the extent to which a focus on simplified food-chains can be unrepresentative of a wider ecological system, as we seek to unpick the effects of seasonal shifts in trophic networks. Perhaps more importantly, those diverse responses illustrate the scope for asynchrony to be buffered within these food-webs: at the level of individual caterpillars and their populations, and also for the secondary consumers that feed on them. Properly characterising the diversity of trophic interactions, the extent to which the impacts of asynchrony vary among those interactions, and the degree to which these systems are inherently resilient to mis-timing is vital—not only for robustly forecasting the ecological impacts of phenological change, but to understand the importance of trophic mismatch as an emergent threat for populations, species, communities and ecosystems in a warming world.

## Acknowledgements

Many of the ideas articulated in this paper arose through years of conversation with colleagues and collaborators, for which we are immensely grateful. Specifically, we would like to mention Kirsty Macphie, Darren Obbard, Megan Stamp, Graham Stone, and Chris Thomas. We would also like to thank Andrew Law (City of Edinburgh Council, Natural Heritage Officer) who kindly gave permission to carry out fieldwork at the Hermitage of Braid LNR, Edinburgh. Jarrod Hadfield (University of Edinburgh) offered helpful statistical advice on an early version of this study.

## Author contributions

JCW conceived the study and carried out the experiment. ABP contributed to conceptual development and statistical analysis, including developing the simulation analysis in the Appendix. JCW led the writing, with both authors contributing substantially to writing and editing.

## Data Availability

Code and data necessary to reproduce the results in this paper are archived at: https://github.com/jamiecweir/Hugo_de_Vries.

## Methods

### Caterpillar species

We selected six phytophagous British macrolepidopteran moth species as our focal taxa: the Winter Moth *Operophtera brumata*, the Mottled Umber *Erannis defoliaria* (both Geometridae), the Gypsy Moth *Lymantria dispar*, the Black Arches *Lymantria monacha*, the Vapourer *Orgyia antiqua*, and the Scarce Vapourer *Orgyia recens* (all Erebidae). The larval stage of all species feed in early spring and are generally widespread and common in Great Britain and Europe^29^. All species overwinter as ova, except the Scarce Vapourer which passes this period as a diapausing first instar larva^49,50^.

Female winter moths were collected over two years (from 25 Nov 2019 to 8 Jan 2020 and 17 Nov 2020 to 27 Dec 2020) in the Hermitage of Braid LNR (Edinburgh) using lobster-pot style trunk traps, modelled on those described by^51^, which intercept females as they ascend trees after emergence in mid-winter^32^. After collection, females were placed individually in 75 × 25mm glass phials with a wad of cotton to act as an egg-laying medium. Females were stored at ~5°C in complete darkness and allowed to lay freely. Approximately one month later, all tubes were examined and the dead females were removed. Ova from a total of 126 females were obtained in the winter of 2019 (used in the 2020 experiment) and from 85 females in the winter of 2020 (used in the 2021 experiment).

For the remaining species, ova (or larvae for Scarce Vapourer) were obtained as livestock from entomologists or entomological supply companies across the United Kingdom. For each species, females were obtained from at least ten distinct, wild-collected broods. Most livestock were sourced solely from a single population, the location of which differed among some species (Table S11).

### Rearing

Ova/larvae of all species were stored at 5°C (temperature ranged from 4-7°C) in a Russell Hobbs RHCLRF17 tabletop refrigerator to prevent early hatch or diapause break. As required for the experiment, ova/larvae were removed from cold storage and placed at room temperature (ranging from 17-21°C) to stimulate egg hatching and emergence from diapause. We manipulated the timing of hatching in 5-day intervals, from perfect synchrony (asynchrony of 0 days) to 65 days asynchrony. To achieve this, we removed a subset of ova/larvae, sampled from across all broods for each species, and allowed them to hatch/emerge as a batch (see Fig 1b for an overview of experimental design). Exposure to relatively high temperatures (i.e., room temperature) helped ensure individuals hatched/became active at the same time, despite inter- and intra-brood variation in the temperature requirements for eclosion ^32,52^.

For each asynchrony treatment, we assigned larvae at random to each host-plant treatment group, and to one of two rearing “cultures”—replicates within each treatment group (i.e., the unique combination of asynchrony treatment, caterpillar species, and host-plant species). “Cultures” comprised ten individuals (see Fig 1), which we first reared in small 75 × 50 × 15mm transparent plastic containers and then, at around the third instar, in larger 175 × 100 × 50mm containers. Rearing containers were lined with white absorbent paper towels. Freshly excised host-plant material was placed in each container and examined daily to check its condition and how much remained. Typically, it was replaced daily, and no less frequently than every second day. Larvae were provided with an excess of host-plant material at all times, such that the quantity of food was never a limiting factor to growth. At the completion of their development, caterpillars pupated freely in the tissue at the base of the container. After all larvae had pupated, excess host-plant material was removed and the containers were stored at room temperature. One month after pupation, pupae were removed and stored outdoors under a canopy at ambient environmental temperature (56.06°N, −3.77°E). Pupae were sexed individually by evaluating shape, size, and the presence of genital and anal pores on the abdomen.

### Asynchrony assays

We aimed to quantify the performance effects of increasing asynchrony across a range of host-by-caterpillar species interactions. We selected eight host-plant species that are common and widespread in Great Britain and known to be used by all of the focal caterpillar species (Robinson *et al*., 2010): Alder *Alnus glutinosa*, Apple *Malus domestica*, Birch *Betula pendula*, Hawthorn *Crataegus monogyna*, Oak *Quercus robur*, Sallow *Salix capraea*, Sycamore *Acer pseudoplatanus*, and Willow *Salix alba*.

We manipulated larval asynchrony between egg hatch (or emergence from diapause) and host-plant bud-burst by staggering the removal of moth eggs from cold storage as the foliage aged naturally in the field. The experiment was initiated for each species when the required number of individual trees could be sampled in the field at the “first full leaf” stage—this was taken as the date at which the first fully unfurled leaf with a recognisable shape (though not full-sized) was observed in the field^cf.7,21^. We designated hatching at the start of the experiment as perfect temporal match (synchrony) with the leafing phenology of the host tree, and hence an asynchrony of zero. The experiment began on different calendar dates for each host-plant species, due to the need to find individual trees of each species at the appropriate phenological stage, though these were co-ordinated to minimise the variation in start date (Table S12), with a maximum difference in start date of 11 days between Hawthorn and Oak in 2021. The experiment was conducted over two years (2020 and 2021) and host-caterpillar treatments differed slightly between years: in 2020, the experiment included only Winter Moth, Mottled Umber, and Vapourer caterpillars, whereas in 2021 (*n* = 4250) the experiment included all caterpillar species on all host-plants (*n* = 13440).

Fresh foliage was collected from at least 12 individual trees of each tree species every two to three days. Cut sprigs 15cm in length were stored in airtight plastic bags at 5°C until required for feeding, to maintain freshness. Foliage was collected from trees near Falkirk (Stirlingshire; 56.069°N, −3.767°E) and Kincardine (Fife; 56.057°N, −3.613°E). Because asynchrony was simulated by staggering caterpillar hatching, all the individuals active at any one time were fed on samples from the same foliage collection. The secondary chemistry of plant leaves can vary not only between species but also between individuals of the same species, due to local differences in environmental conditions, soil chemistry, local geology, and associated plant species (for example^53–55^. This variation could affect the performance of insects feeding on those leaves. To minimise the effects of individual variation in leaf properties *within a species*, leaves from across all the sampled tree individuals were randomly assigned to each rearing culture, such that caterpillars at any one time had access to foliage from a range of different host-plant individuals of the same host-plant species.

Caterpillar performance was quantified by assessing:

i. the **survival** (0/1) of each individual from hatch to pupation.
ii. the final **pupal mass** (mg) attained by each individual one month after pupation, measured using a Mettler AJ50 balance. This is a widely used and robust predictor of adult female fecundity ^33^. Pupae were sexed when weighed.
iii. the **development time** (days) from egg hatch to pupation. A longer developmental period potentially exposes caterpillars to increased predation, parasitism, and exposure to adverse environmental conditions.

Since larvae were reared in groups from multiple females it was not possible to allocate maternal IDs to individuals (see Appendix 1 for a simulation test that examines the impact of this aspect of experimental design on model inferences).

### Statistical analysis

We fitted generalised linear mixed models (GLMMs) to the data in *R* v. 4.4.1 (R Development Core Team, 2004) using the Bayesian package MCMCglmm^56^. To evaluate changes in performance associated with asynchrony, we separately modelled survival to pupation (binomial response), pupal mass (Gaussian response), and development time (Gaussian response). All models included asynchrony in days, year (as a factor), and the day-by-year interaction as fixed effects. As sex differences in caterpillar mass can be large, we included sex and two-way interactions between sex and year and asynchrony in the pupal mass and development time models. To estimate variance in the performance of caterpillars that are synchronous with different host-plants, we included caterpillar species, host-plant species, and a host-caterpillar interaction as random intercept terms. To estimate the extent to which the impacts of asynchrony vary among caterpillar species, host-plant species, and host-by-caterpillar interactions, we also included these terms as random slopes. To accommodate the potential that host-plant material might mature at different rates between the two experimental years (2020 and 2021), all models also allowed for the intercept and asynchrony slope to vary across host-by-year combinations. For pupal mass and development time we also allowed for the intercept and asynchrony slope to vary across unique combinations of (*i*) sex and caterpillar species, (*ii*) sex and host, (*iii*) sex, caterpillar species, and host. Model settings were as follows. Survival model: iterations = 11 million, burn-in = 1 million, posterior samples = 50,000. Pupal mass and duration models: 30 million iterations, burnin = 10 million, 10,000 samples of posterior. All models included a residual term. All fixed and random terms were estimated with an effective sample size >1000.

We used default priors for the fixed effects (mean = 0, with a large variance), inverse Wishart priors on the residuals, and used parameter expanded priors (scaled F_1,1_ distribution, with scale = 1000) for the remaining random effects. Previous studies suggest we might expect a quadratic relationship between performance and degree of asynchrony^7,57^, however we found from inspection of the model fits against the raw data that a simple linear association fit the data well for the average declines arising from hatching after bud-burst (Figure 4). From our models, we generated predictions of the performance effects of asynchrony as follows:

a. averaged across all host-plant and caterpillar species combinations (Fig 1);
b. on each host-plant species, averaged across all caterpillar species (Fig 4a);
c. on each caterpillar species, averaged across all host-plant species (Fig 4b); and,
d. on each host-caterpillar species combination (Fig 4c-h).

To estimate the relative explanatory contribution of our key focal parameters (host-plant species, caterpillar species, and a host-by-caterpillar interaction), we estimated the variance (on latent scale for the survival model) explained by the fixed effects and random effects^58^. We then used this variance decomposition to examine the variance in performance that arose from differences in the intercept (baseline) versus the effect of slope (i.e., impact of asynchrony) and the contributions of among host, caterpillar, and host-caterpillar variation.

