## Supplementary Materials for "The impact of phenological mismatch varies across woodland food-web interactions"

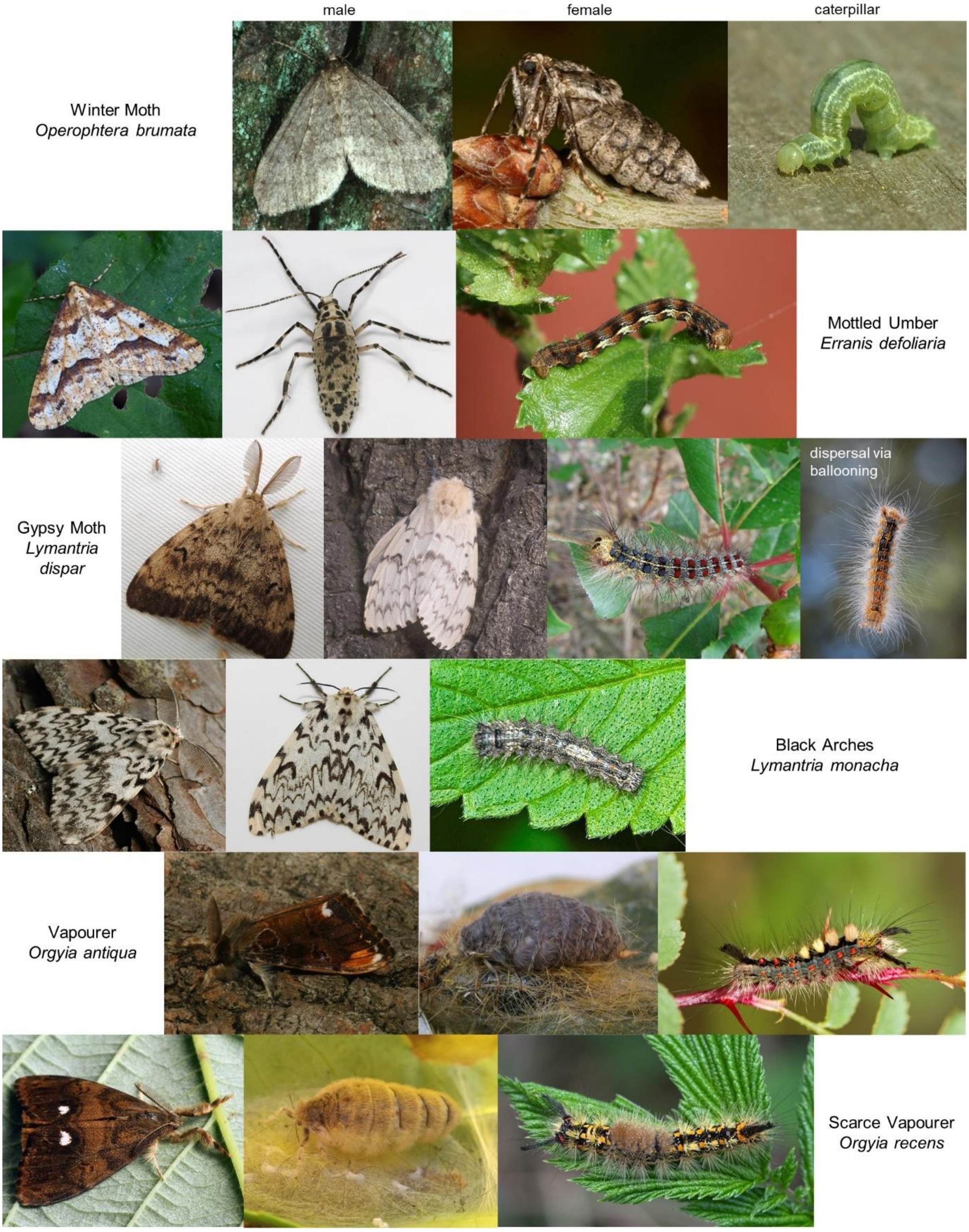

**Fig S1 | Adult moths and caterpillars of study species.** One species per row, with the male, female, and caterpillar illustrated left to right. Gypsy Moth additionally shows a young, early instar caterpillar dispersing to a new location via 'ballooning' on a thread of silk. All images are reproduced under a creative commons licence. Image credits (left to right by row): Louis-Michel Nageleisen, Gyorgy Csoka, Anne Tanne, Urmas Ojango, Janet Graham, S. Rae, Anita Gould, Hedera.Baltica, Wikimedia Commons, Ilia Ustyantsev, Nick Dobbs, Bernd Thaller, Ilia Ustyantsev, Ben Smart, Wikimedia Commons, Ilia Ustyantsev, Will George, Urmas Ojango.

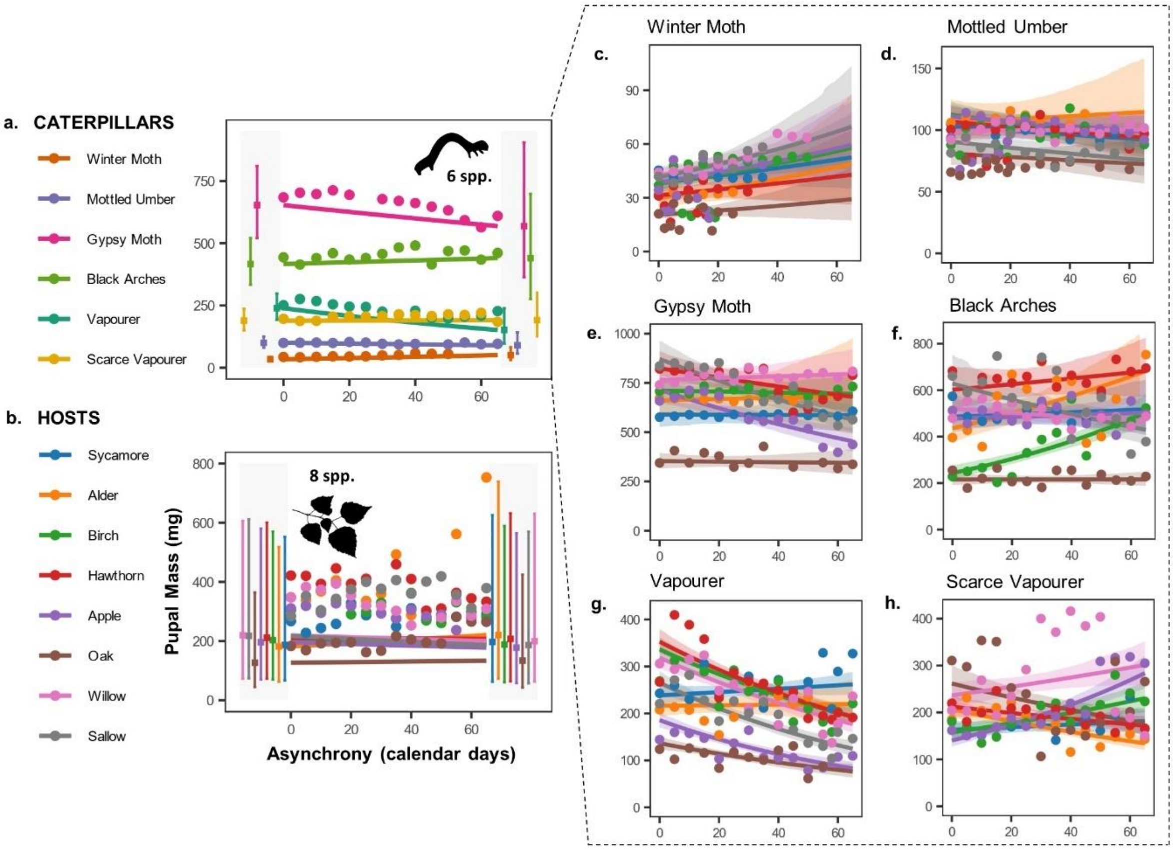

**Fig S2 | Average predicted effects of asynchrony on pupal mass** (mg) across (a) six spring caterpillar species and (b) eight of their host-plant species. Inset panel (c-h) shows the predicted relationships in all pairwise combinations of each caterpillar species fed on each host-plant. Trends represent posterior mean model estimates with shaded 95% credible intervals. Square points in the grey shaded zone of the species average plots (a and b) correspond to model estimates (± 95% CIs) of survival probability at the minimum (0 days) and maximum (65 days) values of induced asynchrony. Round points on all panels show numerical averages of raw data for each level of asynchrony across the relevant groups.

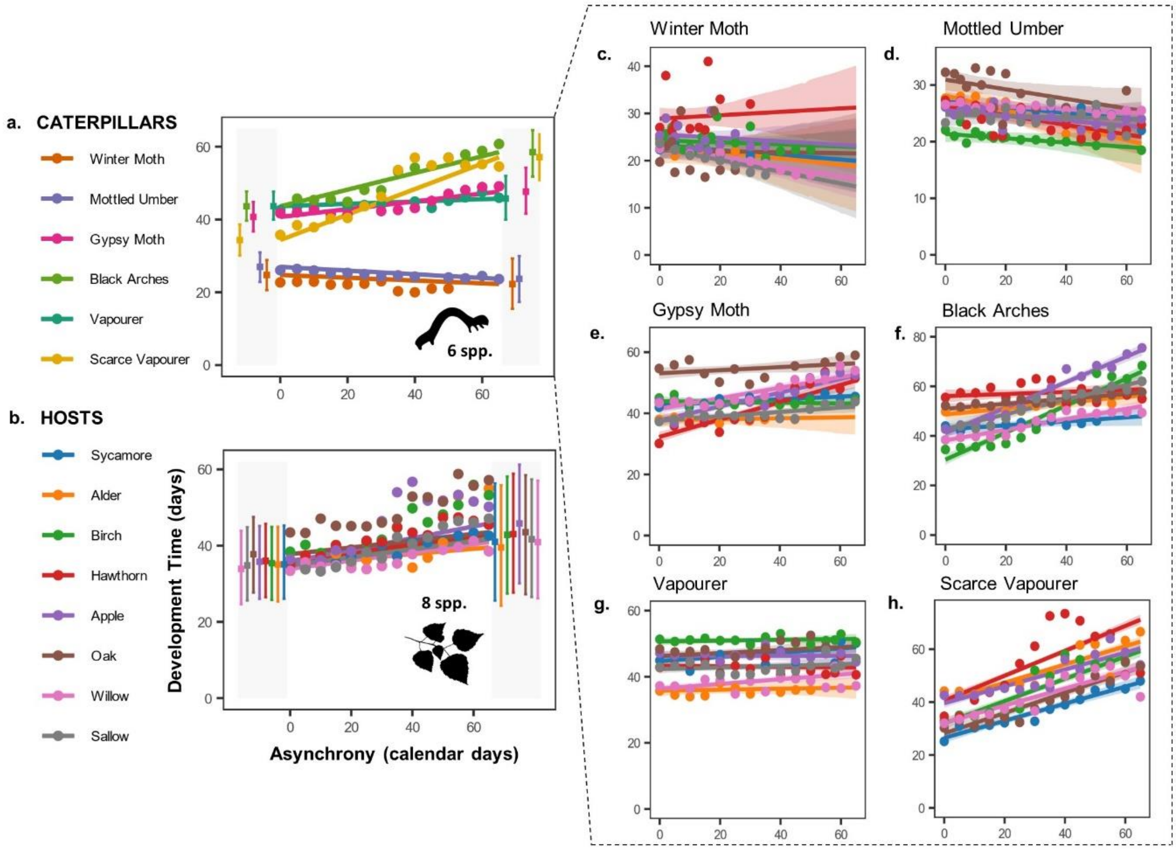

**Fig S3 | Average predicted effects of asynchrony on development time** (calendar days), from egg hatch to pupation, across (a) six spring caterpillar species and (b) eight of their host-plant species. Inset panel (c-h) shows the predicted relationships in all pairwise combinations of each caterpillar species fed on each host-plant. Trends represent posterior mean model estimates with shaded 95% credible intervals. Square points in the grey shaded zone of the species average plots (a and b) correspond to model estimates (± 95% CIs) of survival probability at the minimum (0 days) and maximum (65 days) values of induced asynchrony. Round points on all panels show numerical averages of raw data for each level of asynchrony across the relevant groups.

**Table S1 | GLMM model output: effects of asynchrony on caterpillar survival, across caterpillar and host-plant species.** The experiment was carried out over two years ('Year'), which was fitted as a fixed effect alongside the quantity of Asynchrony (in days). We included random terms to capture the effect of caterpillar species ('Caterpillar'), host-plant species ('Host'), and their interaction ('Caterpillar: Host') on the baseline of individual caterpillar survival (random intercept; i.e., at asynchrony = 0), and on the impact of asynchrony (random slope; i.e., how the Asynchrony effect varies among levels).

| SURVIVAL TIME MODEL | Fixed Effect Coefficient / Random Effect Variance (Mean ± CIs) | Effective sample size | pMCMC |
|---|---|---|---|
| **Fixed Terms** | | | |
| Intercept | -0.26 (-1.72 - 1.22) | 45475 | 0.695 |
| Asynchrony | -0.05 (-0.07 - -0.02) | 48176 | 0.007 |
| Year 2021 | 0.88 (0.28 - 1.49) | 50000 | 0.011 |
| Asynchrony: Year 2021 | -0.005 (-0.015 - 0.005) | 48451 | 0.293 |
| **Random Terms** | | | |
| Culture | 0.016 (0.000 - 0.051) | 34609 | |
| Caterpillar | 2.12 (0.00 - 6.33) | 1825 | |
| Host | 1.11 (0.00 - 3.46) | 3241 | |
| Caterpillar: Host | 1.97 (1.01 - 3.11) | 9213 | |
| Year: Host | 0.31 (0.02 - 0.84) | 11719 | |
| Asynchrony slope: Caterpillar | 0.001 (0.00 - 0.002) | 2624 | |
| Asynchrony slope: Host | 0.0001 (0.0000 - 0.0004) | 13652 | |
| Asynchrony slope: Caterpillar: Host | 0.0004 (0.0001 - 0.0006) | 22675 | |
| Asynchrony slope: Year: Host | 0.0000 (0.0000 - 0.0002) | 27025 | |

**Table S2 | LMM model output: effects of asynchrony on caterpillar pupal mass, across caterpillar and host-plant species.** The experiment was carried out over two years ('Year'), which was fitted as a fixed effect alongside the quantity of Asynchrony (in days). Pupal mass is expected to differ between the sexes, and this was therefore fitted as a fixed effect ('Sex') and also allowed to interact with other key effects. We included random terms to capture the effect of caterpillar species ('Caterpillar'), host-plant species ('Host'), and their interaction ('Caterpillar: Host') on the baseline of individual caterpillar survival (random intercept; i.e., at asynchrony = 0), and on the impact of asynchrony (random slope; i.e., how the Asynchrony effect varies among levels).

| **MASS MODEL** | **Fixed Effect Coefficient / Random Effect Variance (Mean ± CIs)** | **Effective sample size** | **pMCMC** |
|---|---|---|---|
| **Fixed Terms** | | | |
| Intercept | 4.16 (3.11 - 5.22) | 8421 | 0.0006 |
| Asynchrony | -0.006 (-0.014 - 0.002) | 10000 | 0.14 |
| Sex F | 0.84 (0.04 - 1.63) | 10000 | 0.04 |
| Year 2021 | 0.24 (0.09 - 0.40) | 10000 | 0.008 |
| Asynchrony: Year 2021 | 0.005 (-0.001 - 0.012) | 10000 | 0.08 |
| Asynchrony: Sex F | -0.0002 (-0.0037 - 0.0031) | 10000 | 0.87 |
| **Random Terms** | | | |
| Culture | 0.003 (0.002 - 0.004) | 9410 | |
| Caterpillar | 1.23 (0.00 - 3.86) | 732.3 | |
| Host | 0.04 (0.00 - 0.12) | 10000 | |
| Host: Sex | 0.005 (0.000 - 0.002) | 10000 | |
| Caterpillar: Host | 0.008 (0.000 - 0.002) | 10000 | |
| Sex: Caterpillar | 0.44 (0.05 - 1.23) | 3058 | |
| Asynchrony slope: Caterpillar | 0.0000 (0.0000 - 0.0002) | 9891.3 | |
| Asynchrony slope: Host | 0.0000 (0.0000 - 0.0000) | 8396 | |
| Year: Host | 0.023 (0.004 - 0.056) | 10000 | |
| Sex: Caterpillar: Host | 0.002 (0.000 - 0.004) | 10000 | |
| Asynchrony slope: Sex: Caterpillar | 0.0000 (0.0000 - 0.0000) | 10319 | |
| Asynchrony slope: Host: Sex | 0.0000 (0.0000 - 0.0001) | 10000 | |
| Asynchrony slope: Caterpillar: Host | 0.0000 (0.0000 - 0.0001) | 9918 | |
| Asynchrony slope: Year: Host | 0.0000 (0.0000 - 0.0000) | 10000 | |
| Asynchrony slope: Sex: Caterpillar: Host | 0.00002 (0.00000 - 0.00003) | 9036 | |
| *Residual* | 0.068 (0.065 - 0.070) | 10000 | |

**Table S3 | LMM model output: effects of asynchrony on caterpillar development time, across caterpillar and host-plant species.** The experiment was carried out over two years ('Year'), which was fitted as a fixed effect alongside the quantity of Asynchrony (in days). Pupal mass is expected to differ between the sexes, and this was therefore fitted as a fixed effect ('Sex') and also allowed to interact with other key effects. We included random terms to capture the effect of caterpillar species ('Caterpillar'), host-plant species ('Host'), and their interaction ('Caterpillar: Host') on the baseline of individual caterpillar survival (random intercept; i.e., at asynchrony = 0), and on the impact of asynchrony (random slope; i.e., how the Asynchrony effect varies among levels).

| DEVELOPMENT TIME MODEL | Fixed Effect Coefficient / Random Effect Variance (Mean ± CIs) | ESS | pMCMC |
|---|---|---|---|
| **Fixed Terms** | | | |
| Intercept | 29.8 (20.5 - 39.3) | 9455 | 0.0002 |
| Asynchrony | 0.03 (-0.15 - 0.21) | 10000 | 0.66 |
| Sex F | 6.34 (-0.36 - 12.99) | 12055 | 0.06 |
| Year 2021 | -0.56 (-1.32 - 0.20) | 10000 | 0.13 |
| Asynchrony: Year 2021 | 0.05 (0.02 - 0.09) | 10000 | 0.01 |
| Asynchrony: Sex F | 0.02 (-0.07 - 0.10) | 10000 | 0.58 |
| **Random Terms** | | | |
| Culture | 1.86 (1.45 - 2.27) | 10000 | |
| Caterpillar | 91.63 (0.00 - 285.43) | 1924 | |
| Host | 4.96 (0.00 - 19.19) | 7551 | |
| Host: Sex | 1.69 (0.00 - 6.02) | 9457 | |
| Caterpillar: Host | 19.46 (8.73 - 31.28) | 7382 | |
| Sex: Caterpillar | 31.43 (2.25 - 90.59) | 3132 | |
| Asynchrony slope: Caterpillar | 0.05 (0.00 - 0.14) | 3338 | |
| Asynchrony slope: Host | 0.002 (0.00 - 0.057) | 10000 | |
| Year: Host | 0.25 (0.00 - 0.90) | 10000 | |
| Sex: Caterpillar: Host | 7.86 (4.19 - 12.23) | 8677 | |
| Asynchrony slope: Sex: Caterpillar | 0.004 (0.00 - 0.02) | 7270 | |
| Asynchrony slope: Host: Sex | 0.0005 (0.00 - 0.002) | 10000 | |
| Asynchrony slope: Caterpillar: Host | 0.007 (0.003 - 0.011) | 8300 | |
| Asynchrony slope: Year: Host | 0.001 (0.000 - 0.002) | 10000 | |
| Asynchrony slope: Sex: Caterpillar: Host | 0.002 (0.001 - 0.004) | 10297 | |
| *Residual* | 15.83 (15.27 - 16.44) | 10000 | |

**Table S4 | Variance in caterpillar performance and the effects of asynchrony attributable to differences in the resource-consumer species interaction.** Total variance explained by each model (survival, pupal mass, and development time) is partitioned based on that attributed to the fixed effects, variation in the random intercept, and variation in the random slope. Variation explained by the random intercepts and random slopes are further subdivided into the proportion explained by caterpillar (consumer) species ('Caterpillar'), host-plant (resource) species ('Host-plant'), and their interaction ('Interaction'). For example, where the performance effect of asynchrony varies mostly among caterpillar taxa, and the host-plant species a caterpillar is on has minimal impact, we would expect the caterpillar slope to explain a greater proportion of the variance than either the host-plant slope or the interaction slope. If the effect of increasing asynchrony on caterpillar performance varies idiosyncratically among unique resource-consumer species pairings, we would expect the interaction slope to explain a greater proportion of the available variance. Variation in the intercept describes variability in the underlying, baseline performance of a caterpillar at perfect temporal synchrony (i.e., when asynchrony is equal to zero).

| | | Performance Metric | | |
|---|---|---|---|---|
| | **Model Term** | **Survival** | **Mass** | **Development Time** |
| | Total Variance | 8.16 (4.79 - 15.57) | 1.63 (0.66 - 4.63) | 228.17 (107.49 - 523.25) |
| **Total Variance Decomposed** | Fixed | 0.12 (0.02 - 0.23) | 0.12 (0.01 - 0.32) | 0.08 (0.00 - 0.20) |
| | Random intercept | 0.71 (0.47 - 0.90) | 0.78 (0.54 - 0.96) | 0.66 (0.35 - 0.91) |
| | Random slope | 0.16 (0.04 - 0.38) | 0.08 (0.01 - 0.21) | 0.24 (0.04 - 0.54) |
| **Random Variance Decomposed** | Caterpillar intercept | 0.22 (0.00 - 0.61) | 0.58 (0.00 - 0.89) | 0.44 (0.00 - 0.80) |
| | Caterpillar slope | 0.46 (0.09 - 0.87) | 0.27 (0.00 - 0.68) | 0.65 (0.24 - 0.97) |
| | Host-plant intercept | 0.13 (0.00 - 0.44) | 0.02 (0.00 - 0.11) | 0.01 (0.00 - 0.13) |
| | Host-plant slope | 0.06 (0.00 - 0.34) | 0.06 (0.00 - 0.42) | 0.01 (0.00 - 0.13) |
| | Interaction intercept | 0.34 (0.11 - 0.60) | 0.01 (0.00 - 0.02) | 0.13 (0.02 - 0.26) |
| | Interaction slope | 0.38 (0.07 - 0.74) | 0.03 (0.00 - 0.13) | 0.14 (0.01 - 0.32) |

**Table S5 | Model estimate of slope of survival (log odds/day) with increasing asynchrony** in each caterpillar species (on the average host-plant) and on each host-plant species (with the average caterpillar). Model predictions are estimated for 2021, with 95% credible intervals.

| **Caterpillar Species** | | **Host-plant Species** | |
|---|---|---|---|
| Winter Moth | -0.078 (-0.098 - -0.057) | Sycamore | -0.049 (-0.076 - -0.022) |
| Mottled Umber | -0.047 (-0.064 - -0.03) | Alder | -0.059 (-0.089 - -0.031) |
| Gypsy Moth | -0.059 (-0.076 - -0.042) | Birch | -0.048 (-0.074 - -0.021) |
| Black Arches | -0.033 (-0.05 - -0.015) | Hawthorn | -0.049 (-0.075 - -0.021) |
| Vapourer | -0.052 (-0.069 - -0.036) | Apple | -0.054 (-0.081 - -0.028) |
| Scarce Vapourer | -0.04 (-0.058 - -0.023) | Oak | -0.059 (-0.087 - -0.031) |
| | | White Willow | -0.047 (-0.075 - -0.02) |
| | | Sallow | -0.048 (-0.075 - -0.021) |

**Table S6 | Model estimate of slope of change in survival probability (log odds/day) with increasing asynchrony** for each host-caterpillar species pairing. Model predictions are estimated for year 2021 and are shown with 95% credible intervals.

| Host-plant Species | Caterpillar Species | | | | | |
|---|---|---|---|---|---|---|
| | Winter Moth | Mottled Umber | Gypsy Moth | Black Arches | Vapourer | Scarce Vapourer |
| Sycamore | -0.087 | -0.049 | -0.042 | -0.047 | -0.014 | -0.039 |
| | (-0.115 - -0.060) | (-0.064 - -0.034) | (-0.058 - -0.025) | (-0.065 - -0.029) | (-0.028 - 0.000) | (-0.053 - -0.024) |
| Alder | -0.084 | -0.060 | -0.080 | -0.048 | -0.065 | -0.060 |
| | (-0.121 - -0.047) | (-0.084 - -0.038) | (-0.105 - -0.057) | (-0.068 - -0.030) | (-0.08 - -0.050) | (-0.101 - -0.018) |
| Birch | -0.060 | -0.053 | -0.074 | -0.013 | -0.022 | -0.046 |
| | (-0.078 - -0.042) | (-0.068 - -0.038) | (-0.090 - -0.057) | (-0.027 - 0.000) | (-0.037 - -0.006) | (-0.064 - -0.029) |
| Hawthorn | -0.087 | -0.040 | -0.061 | -0.015 | -0.064 | -0.016 |
| | (-0.117 - -0.058) | (-0.057 - -0.024) | (-0.077 - -0.045) | (-0.032 - 0.002) | (-0.079 - -0.050) | (-0.030 - -0.002) |
| Apple | -0.092 | -0.041 | -0.062 | -0.050 | -0.053 | -0.041 |
| | (-0.113 - -0.071) | (-0.054 - -0.028) | (-0.078 - -0.046) | (-0.065 - -0.035) | (-0.066 - -0.041) | (-0.056 - -0.027) |
| Oak | -0.094 | -0.055 | -0.064 | -0.038 | -0.066 | -0.070 |
| | (-0.122 - -0.065) | (-0.075 - -0.035) | (-0.083 - -0.047) | (-0.053 - -0.023) | (-0.081 - -0.052) | (-0.089 - -0.052) |
| White Willow | -0.053 | -0.019 | -0.078 | -0.021 | -0.072 | -0.010 |
| | (-0.073 - -0.032) | (-0.034 - -0.005) | (-0.095 - -0.061) | (-0.035 - -0.007) | (-0.088 - -0.057) | (-0.024 - 0.005) |
| Sallow | -0.096 | -0.050 | -0.023 | -0.009 | -0.058 | -0.025 |
| | (-0.121 - -0.071) | (-0.066 - -0.034) | (-0.038 - -0.010) | (-0.025 - 0.007) | (-0.071 - -0.045) | (-0.039 - -0.012) |

**Table S7 | Model estimate of the slope of change in final pupal mass (mg/day) with increasing asynchrony** in each caterpillar species (on the average host-plant) and on each host-plant species (with the average caterpillar). Model predictions are estimated for females and the year 2021, with 95% credible intervals. Pupal mass is a reliable proxy of female fecundity in many spring caterpillar species (see Weir. 2024. *Oecologia*), and is therefore a useful metric of performance (i.e., reproductive success).

| **Caterpillar Species** | | **Host-plant Species** | |
|---|---|---|---|
| Winter Moth | 0.006 (-0.001 - 0.013) | Sycamore | 0.001 (-0.006 - 0.008) |
| Mottled Umber | -0.002 (-0.008 - 0.005) | Alder | 0.003 (-0.004 - 0.010) |
| Gypsy Moth | -0.002 (-0.008 - 0.004) | Birch | -0.001 (-0.008 - 0.006) |
| Black Arches | 0.001 (-0.006 - 0.007) | Hawthorn | 0.000 (-0.007 - 0.007) |
| Vapourer | -0.007 (-0.013 - -0.001) | Apple | -0.001 (-0.008 - 0.005) |
| Scarce Vapourer | 0.000 (-0.006 - 0.007) | Oak | 0.001 (-0.006 - 0.008) |
| | | White Willow | -0.001 (-0.008 - 0.005) |
| | | Sallow | -0.002 (-0.009 - 0.005) |

**Table S8 | Model estimate of slope of change in pupal mass (mg/day) with increasing asynchrony** for each host-caterpillar species pairing. Model predictions are estimated for females and the year 2021, and are shown with 95% credible intervals. Pupal mass is a reliable proxy of female fecundity in many spring caterpillar species (see Weir. 2024. *Oecologia*), and is therefore a useful metric of performance (i.e., reproductive success).

| **Host-plant Species** | **Caterpillar Species** | | | | | |
|---|---|---|---|---|---|---|
| | Winter Moth | Mottled Umber | Gypsy Moth | Black Arches | Vapourer | Scarce Vapourer |
| Sycamore | 0.005<br>(-0.003 - 0.012) | -0.002<br>(-0.005 - 0.001) | 0.000<br>(-0.004 - 0.004) | 0.001<br>(-0.004 - 0.006) | 0.001<br>(-0.001 - 0.004) | 0.002<br>(-0.002 - 0.005) |
| Alder | 0.008<br>(-0.002 - 0.018) | 0.001<br>(-0.005 - 0.008) | 0.001<br>(-0.006 - 0.007) | 0.007<br>(0.002 - 0.012) | 0.000<br>(-0.004 - 0.005) | 0.004<br>(-0.002 - 0.009) |
| Birch | 0.007<br>(0.002 - 0.011) | -0.003<br>(-0.007 - 0.001) | 0.000<br>(-0.003 - 0.002) | 0.011<br>(0.008 - 0.015) | -0.009<br>(-0.011 - -0.007) | -0.007<br>(-0.009 - -0.004) |
| Hawthorn | 0.005<br>(-0.004 - 0.013) | -0.002<br>(-0.006 - 0.002) | -0.003<br>(-0.006 - 0.000) | 0.002<br>(-0.002 - 0.006) | -0.009<br>(-0.012 - -0.007) | 0.006<br>(0.002 - 0.009) |
| Apple | 0.007<br>(0.000 - 0.013) | -0.002<br>(-0.005 - 0.001) | -0.007<br>(-0.010 - -0.004) | 0.001<br>(-0.003 - 0.004) | -0.012<br>(-0.016 - -0.009) | -0.003<br>(-0.006 - 0.000) |
| Oak | 0.005<br>(-0.003 - 0.014) | -0.002<br>(-0.007 - 0.003) | 0.000<br>(-0.004 - 0.003) | 0.000<br>(-0.003 - 0.003) | -0.009<br>(-0.013 - -0.005) | 0.011<br>(0.008 - 0.014) |
| White Willow | 0.007<br>(0.002 - 0.013) | 0.000<br>(-0.002 - 0.003) | 0.001<br>(-0.002 - 0.004) | -0.002<br>(-0.005 - 0.001) | -0.009<br>(-0.011 - -0.007) | -0.006<br>(-0.010 - -0.002) |
| Sallow | 0.008<br>(0.001 - 0.015) | -0.003<br>(-0.007 - 0.001) | -0.007<br>(-0.009 - -0.004) | -0.006<br>(-0.010 - -0.002) | -0.012<br>(-0.014 - -0.009) | 0.004<br>(0.000 - 0.007) |

**Table S9 | Model estimate of the slope of change in larval development time (days/day) with increasing asynchrony** for each caterpillar species (on the average host-plant) and on each host-plant species (with the average caterpillar). Model predictions are estimated for females and the year 2021, with 95% credible intervals.

| **Caterpillar Species** | | **Host-plant Species** | |
|---|---|---|---|
| Winter Moth | -0.038 (-0.134 - 0.05) | Sycamore | 0.091 (-0.092 - 0.282) |
| Mottled Umber | -0.051 (-0.129 - 0.025) | Alder | 0.069 (-0.123 - 0.262) |
| Gypsy Moth | 0.107 (0.030 - 0.183) | Birch | 0.114 (-0.071 - 0.308) |
| Black Arches | 0.229 (0.154 - 0.313) | Hawthorn | 0.108 (-0.093 - 0.283) |
| Vapourer | 0.032 (-0.043 - 0.106) | Apple | 0.154 (-0.042 - 0.340) |
| Scarce Vapourer | 0.350 (0.269 - 0.432) | Oak | 0.089 (-0.107 - 0.272) |
| | | White Willow | 0.108 (-0.068 - 0.309) |
| | | Sallow | 0.106 (-0.082 - 0.289) |

**Table S10 | Model estimate of slope of change in larval development time (days/day) with increasing asynchrony** for each host-caterpillar species pairing. Model predictions are estimated for females and the year 2021, and are shown with 95% credible intervals.

| **Host-plant Species** | **Caterpillar Species** | | | | | |
|---|---|---|---|---|---|---|
| | Winter Moth | Mottled Umber | Gypsy Moth | Black Arches | Vapourer | Scarce Vapourer |
| Sycamore | -0.050<br>(-0.179 - 0.074) | -0.050<br>(-0.100 - 0.003) | 0.046<br>(-0.016 - 0.107) | 0.082<br>(-0.002 - 0.166) | 0.064<br>(0.028 - 0.100) | 0.323<br>(0.268 - 0.381) |
| Alder | -0.066<br>(-0.247 - 0.111) | -0.131<br>(-0.243 - -0.021) | 0.015<br>(-0.092 - 0.128) | 0.120<br>(0.035 - 0.194) | 0.015<br>(-0.052 - 0.080) | 0.315<br>(0.210 - 0.416) |
| Birch | -0.021<br>(-0.092 - 0.044) | -0.038<br>(-0.094 - 0.022) | -0.012<br>(-0.061 - 0.038) | 0.547<br>(0.489 - 0.607) | 0.012<br>(-0.021 - 0.045) | 0.346<br>(0.300 - 0.390) |
| Hawthorn | 0.036<br>(-0.112 - 0.184) | -0.085<br>(-0.15 - -0.018) | 0.283<br>(0.230 - 0.336) | 0.042<br>(-0.032 - 0.115) | -0.007<br>(-0.043 - 0.027) | 0.421<br>(0.366 - 0.474) |
| Apple | -0.037<br>(-0.146 - 0.062) | -0.045<br>(-0.095 - 0.003) | 0.226<br>(0.178 - 0.276) | 0.515<br>(0.459 - 0.569) | 0.000<br>(-0.051 - 0.05) | 0.471<br>(0.422 - 0.519) |
| Oak | -0.010<br>(-0.154 - 0.13) | -0.085<br>(-0.165 - -0.005) | 0.050<br>(-0.015 - 0.112) | 0.100<br>(0.045 - 0.156) | 0.045<br>(-0.019 - 0.104) | 0.319<br>(0.271 - 0.367) |
| White Willow | -0.099<br>(-0.183 - -0.014) | -0.026<br>(-0.07 - 0.019) | 0.172<br>(0.120 - 0.223) | 0.213<br>(0.163 - 0.264) | 0.072<br>(0.035 - 0.109) | 0.385<br>(0.324 - 0.448) |
| Sallow | -0.125<br>(-0.247 - -0.002) | -0.012<br>(-0.074 - 0.058) | 0.072<br>(0.024 - 0.121) | 0.296<br>(0.219 - 0.370) | 0.025<br>(-0.019 - 0.070) | 0.321<br>(0.260 - 0.379) |

**Table S11 | Source populations of caterpillar livestock.** The same populations were used across each year (2020 and 2021), though the number of source females varied. Caterpillars from at least ten distinct broods were used in each case.

| **Species** | **Source Population** | **Year** | **Brood Number** |
|---|---|---|---|
| Winter Moth<br>*Operophtera brumata* | Edinburgh<br>(55.9°N, -3.2°E) | 2019/20<br>2020/21 | 126<br>84 |
| Mottled Umber<br>*Erannis defoliaria* | Edinburgh (55.9°N, -3.2°E)<br>*and*<br>Callander (56.2°N, -4.2°E) | 2019/20<br>2020/21 | 18<br>24 |
| Gypsy Moth<br>*Lymantria dispar* | Isle of Man (54.2°N, -4.5°E) | 2020/21 | 25 |
| Black Arches<br>*Lymantria monacha* | Newcastle (55.0°N, -1.6°E) | 2020/21 | 25 |
| Vapourer<br>*Orgyia antiqua* | Edinburgh (55.9°N, -3.2°E) | 2019/20<br>2020/21 | 30<br>45 |
| Scarce Vapourer<br>*Orgyia recens* | Bristol (51.4°N, -2.5°E) | 2020/21 | 16 |

**Table S12 | Start dates of the asynchrony experiment on different host-plants, in 2020 and 2021.** The leafing phenology of each tree species varies in the field. For each resource-consumer treatment group, an asynchrony of zero days (i.e., perfect temporal synchrony) was taken as the timing of the first recognisable leaf breaking (see Methods). The treatment groups for each host-plant species therefore began on different calendar dates, as indicated below. Variation in the calendar date timing of each treatment group was minimised as far as possible, but was limited based on the observed phenology of each species in the field.

| | **Year** | |
|---|---|---|
| **Host-plant Species** | **2020** | **2021** |
| Sycamore | 1 May | 13 May |
| Alder | 29 Apr | 15 May |
| Birch | 1 May | 12 May |
| Hawthorn | 29 Apr | 5 May |
| Apple | 29 Apr | 12 May |
| White Willow | 29 Apr | 11 May |
| Sallow | 2 May | 13 May |
| Oak | 4 May | 16 May |
