## Appendix for "The impact of phenological mismatch varies across woodland food-web interactions"

### Simulations to assess the effects of mixed-brood rearing cultures on accuracy, precision, power, and false positive rate.

Appendix to Weir, J. C. and Phillimore, A. B. (2026). The impact of phenological mismatch varies across woodland food-web interactions. *bioRxiv*.

#### Introduction

In total we reared 18,720 caterpillars and for logistical reasons individuals were reared in 'cultures' of ten individuals. Cultures were composed of individuals randomly allocated from different mothers. As a result, we cannot associate individuals with a maternal identity, and this aspect of the experimental design has the potential to introduce pseudoreplication that is unaccounted for in our statistical model and which could impact on our inferences. We anticipate that the impact of such pseudoreplication on inferences will be greater when the among-brood (maternal) variance in offspring performance is large relative to the among-individual variance (see Weir 2024). Earlier work examining the survival and growth of Winter Moth (*Operophtera brumata*) caterpillars found that the among-individual variance was approximately an order of magnitude greater than the among brood variance (Weir 2023). If the same pattern holds across other moth species, then we anticipate the unaccounted for psuedoreplication due to brood variance will be quite minor.

To examine the sensitivity of key focal estimates to different levels of among-individual and among-brood variances we simulate datasets with the same design as our own and then examine the ability of our modelling approach to recover the true main effect for the slope and the true variance values. We use this approach to estimate bias, precision, power, and the false positive ('type I error') rate for each focal variable and to assess the sensitivity of these properties to different levels of among-individual (i.e., within-brood) and among-brood variance. Our focus is on estimating the main effect for the slope (the overall average effect of asynchrony) and the intercept and slope variances among (*i*) caterpillar species, (*ii*) host-plant species, and (*iii*) host-by-caterpillar combinations While we do present additional model outputs in our study, their behaviour will be largely governed by the focal parameters that we consider here.

#### Methods

##### *Simulation structure*

We designed simulations such that they retain the broad structure, sample sizes and sampling procedures as the experiment, with six caterpillar species, eight host plants, 248 broods (reflecting the differences in brood number across the different caterpillar species: e.g., 126 for winter moth), with two cultures per host-by-caterpillar combination at each asynchrony level (0 to 65 days in 5-day increments, totalling 1344 cultures) for a total of 13440 individuals. Individuals from a brood could be selected and assigned at random across different host taxa and across different asynchrony treatments. Our aim with the simulations was to gain broad insight into the extent to which the main focal parameters can be estimated robustly. As our interest is in diagnosing the impact of the study design on our general ability to draw robust inferences, for simplicity, we limit our analysis to the data collected in 2021, do not consider sex, and only consider a Gaussian response.

For each replicate the simulations were set up as follows:

The global intercept was 100, with an average asynchrony slope of -1. We assigned random normal intercept deviations (mean = 0) to each caterpillar species, host-plant species, host-by-caterpillar combination, brood, culture, and individual (see Table A1 for variances). The sum of the global intercept and these random intercept deviations meant that all individuals had a unique intercept value.

We assigned random normal slope deviations (mean = 0) to the slope for each caterpillar species, host species, host-by-caterpillar combination, brood, and individual (Table A1). The sum of the asynchrony slope and the slope deviations meant that all individuals had a unique slope value. The Gaussian response for an individual given a particular level of asynchrony was then calculated as the individual's intercept plus the product of the individual's asynchrony slope and asynchrony treatment.

Our simulations considered high and low values of individual and brood intercept (high = 5, low = 0.5) and slope (high = 0.0625, low = 0.00625) variances. This design allowed us to examine whether brood variance that is unaccounted for in our model design poses an issue for inference. Under the high variance scenarios, the brood and individual variances are half the magnitude of the variances that we used for caterpillar, host, and host-by-caterpillar. Low variance was generated as 0.1 times the value of the high variance scenario.

Our simulations capture hypotheses under which (*i*) there is non-zero variance in intercept and slope among caterpillar species, host species, and their interaction, and (*ii*) the expectation, where the simulated variance for a term is 0. We assume that the among-culture variance is small, consistent with what we observed in this study and in earlier work (Weir 2024).

##### *Parameter Estimation*

For each simulated dataset we estimated parameters using a linear mixed model (LMM) that included a fixed effect of asynchrony, with caterpillar species, host, host-by-caterpillar, culture, and a residual as random intercept terms, and used random slope terms to allow the effect of asynchrony to vary across caterpillar species, hosts, and host-by-caterpillar combinations. The model was fitted using MCMCglmm

(Hadfield 2010). The model structure is equivalent to our survival model, except for having a different response family. For each random term we used the median of the posterior distributions as our parameter estimate. To assess the significance of variance estimates for the focal random terms we obtain the proportion of the posterior that is < 0.01; where this proportion is < 0.05 we infer that the parameter estimate departs from 0 (i.e., is significant).

For each parameter combination (Table A1) we conducted 100 simulations. To assess accuracy and precision of model estimates we calculated the median and 95% quantiles of parameter estimates across the 100 simulations. We infer that an estimate is accurate where the median does not depart substantially from the true simulated value, and we infer the estimate is precise if the 95% quantile of estimated values is tightly distributed around the true value. To assess the power to detect each focal effect, we quantified the proportion of simulations in which a focal term was significant ($p < 0.05$). Finally, we assessed the false positive rate as the proportion of simulations for which a term was significant when the true simulated effect was 0.

For each simulated dataset, we ran MCMCglmm for 800,000 iterations, with the first 100,000 iterations removed as burn-in, resulting in 35,000 samples of the posterior. Default MCMCglmm priors were used for the fixed effects, with inverse Wishart priors used for the residual term, and parameter expanded priors used for the remaining random terms, as described in the Methods. Across the 100 simulations for each parameter combination the median effective sample size of the least well-estimated parameter was > 100, and for 12 out of 16 parameter combinations this value was > 200.

#### Results

##### *Slope main effect estimate*

The main slope effect was estimated precisely and accurately across simulations, regardless of levels of among-individual and among-brood variance (Fig A1D).

##### *Intercept variance*

The among-caterpillar species intercept variance was generally estimated with little bias (Fig A1A), with both bias and precision appearing to be insensitive to the amount of among-individual and among-brood variance. When the among caterpillar intercept variance was simulated to be 0, models correctly estimated the variance to be low. However, across simulation scenarios, the false positive rate was substantially elevated, with this issue most severe when the among brood variance was high (Fig A1A).

The among-host-plant intercept variance followed a very similar pattern to that described above for caterpillar species, with variance estimates showing little evidence of bias (Fig A1B). For this variable too, there was evidence of a substantially elevated false positive rate, with the issue most pronounced under scenarios with high brood variance (Fig A1B).

The host-by-caterpillar interaction intercept variance was found to be accurate and more precise than for the among-caterpillar and/or among-host variance terms (Fig A1C). We found no evidence for an elevated false positive rate in this variable (Fig A1C).

Power was found to be high across all simulations.

##### *Slope variance*

The true slope variance term was estimated accurately across all scenarios and for caterpillar species (Fig A1E), host species (Fig A2F), and the host-by-caterpillar interaction (Fig A2G), with good precision in all cases, but greatest precision for the interaction (Fig A2G). The false positive rate was below 0.05 in all cases.

#### Conclusions

Our simulations demonstrated that most of the parameters that are of interest in our study can be recovered accurately, precisely, and with a false positive rate below 0.05. We identified that there was no issue in the estimation of the three slope variances or for the host-by-caterpillar intercept variance, with all of these variables still behaving well even when the among-brood variance was high.

For the among-caterpillar and among-host intercept variance terms, we estimate an elevated false positive rate, though when the true variance for these terms was set at 0 then the estimated values were also low. High levels of among-brood variance were associated with a further elevation of the false positive rate for these variables.

On this basis of our simulations, we advise caution in interpreting the variance estimates for the caterpillar and host-plant intercept terms, especially where the variance estimate is close to zero. Consequently, some care must be taken when interpreting small magnitude differences in intercept between caterpillar species and host-plant species.

**Table A1 | Variances values used in simulations.** Focal hypothesis (*a*, *b*, *c*, *d*) and individual and brood variance (high = H, low = L) scenarios match terminology in Figure A1.

| Focal hypotheses (which terms have zero variance) | Individual (I) and Brood (B) variance | Caterpillar species intercept variance | Host intercept variance | Host by Caterpillar intercept variance | Brood intercept variance | Culture intercept variance | Individual intercept variance | Caterpillar species slope variance | Host slope variance | Host by Caterpillar slope variance | Brood slope variance | Individual slope variance |
|---|---|---|---|---|---|---|---|---|---|---|---|---|
| *a* = none | IH:BL | 10 | 10 | 10 | 0.5 | 1 | 5 | 0.125 | 0.125 | 0.125 | 0.00625 | 0.0625 |
| *a* = none | IH:BH | 10 | 10 | 10 | 5 | 1 | 5 | 0.125 | 0.125 | 0.125 | 0.0625 | 0.0625 |
| *b* = caterpillar | IH:BL | 0 | 10 | 10 | 0.5 | 1 | 5 | 0 | 0.125 | 0.125 | 0. 00625 | 0.0625 |
| *b* = caterpillar | IH:BH | 0 | 10 | 10 | 5 | 1 | 5 | 0 | 0.125 | 0.125 | 0.0625 | 0.0625 |
| *c* = host-plant | IH:BL | 10 | 0 | 10 | 0.5 | 1 | 5 | 0.125 | 0 | 0.125 | 0. 00625 | 0.0625 |
| *c* = host-plant | IH:BH | 10 | 0 | 10 | 5 | 1 | 5 | 0.125 | 0 | 0.125 | 0.0625 | 0.0625 |
| *d* = interact. | IH:BL | 10 | 10 | 0 | 0.5 | 1 | 5 | 0.125 | 0.125 | 0 | 0. 00625 | 0.0625 |
| *d* = interact. | IH:BH | 10 | 10 | 0 | 5 | 1 | 5 | 0.125 | 0.125 | 0 | 0.0625 | 0.0625 |
| *a* = none | IL:BL | 10 | 10 | 10 | 0.5 | 1 | 0.5 | 0.125 | 0.125 | 0.125 | 0. 00625 | 0.00625 |
| *a* = none | IL:BH | 10 | 10 | 10 | 5 | 1 | 0.5 | 0.125 | 0.125 | 0.125 | 0.0625 | 0.00625 |
| *b* = caterpillar | IL:BL | 0 | 10 | 10 | 0.5 | 1 | 0.5 | 0 | 0.125 | 0.125 | 0. 00625 | 0.00625 |
| *b* = caterpillar | IL:BH | 0 | 10 | 10 | 5 | 1 | 0.5 | 0 | 0.125 | 0.125 | 0.0625 | 0.00625 |
| *c* = host-plant | IL:BL | 10 | 0 | 10 | 0.5 | 1 | 0.5 | 0.125 | 0 | 0.125 | 0. 00625 | 0.00625 |
| *c* = host-plant | IL:BH | 10 | 0 | 10 | 5 | 1 | 0.5 | 0.125 | 0 | 0.125 | 0.0625 | 0.00625 |
| *d* = interact. | IL:BL | 10 | 10 | 0 | 0.5 | 1 | 0.5 | 0.125 | 0.125 | 0 | 0. 00625 | 0.00625 |
| *d* = interact. | IL:BH | 10 | 10 | 0 | 5 | 1 | 0.5 | 0.125 | 0.125 | 0 | 0.0625 | 0.00625 |

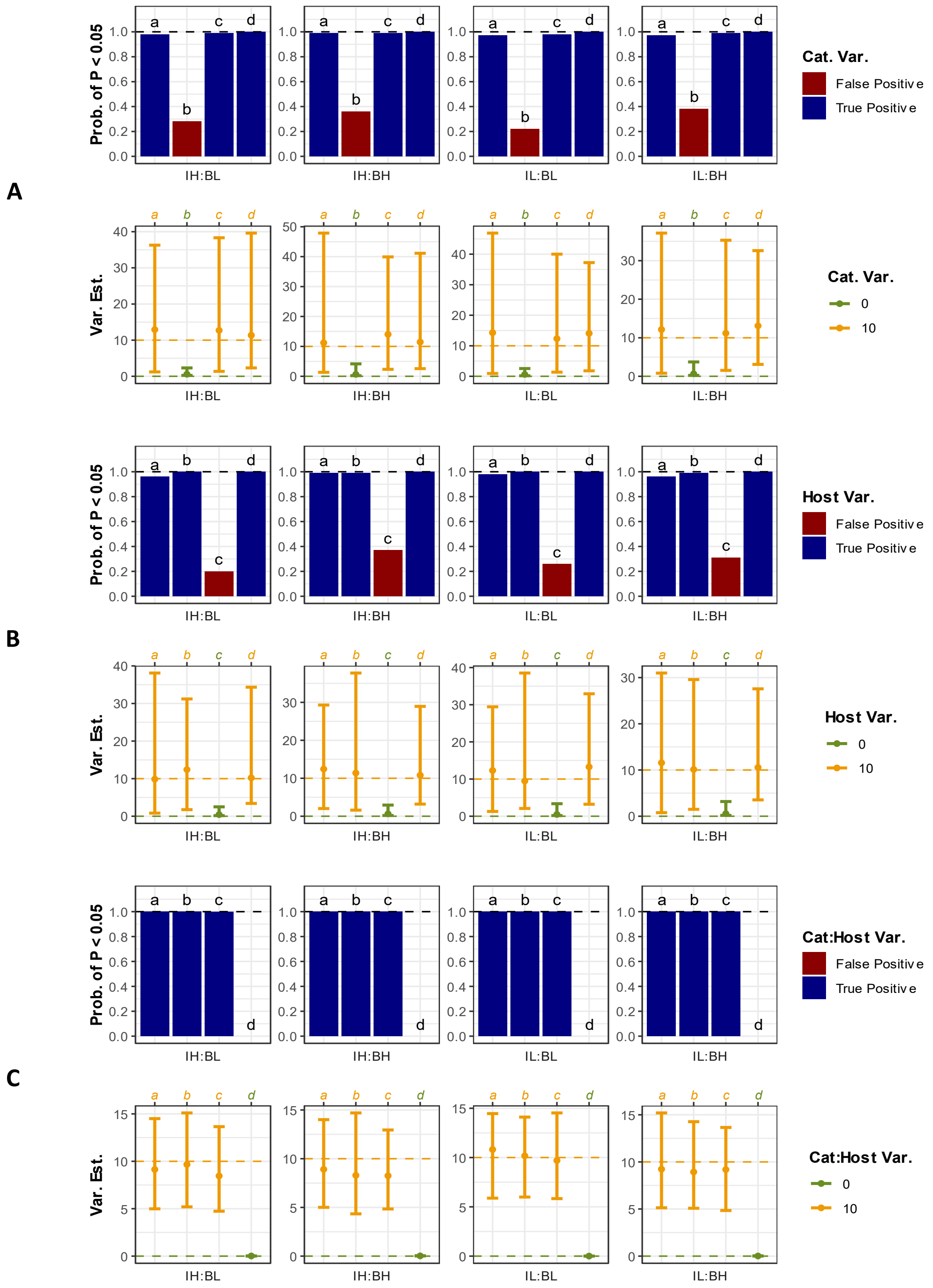


**Fig A1 | Summary of power and false positive rate** (bar plots), and the accuracy and precision estimates (estimation plots) assessing different hypotheses (*a*, *b*, *c*, *d* – for details see Table A1) for the random intercept variance estimates (A-C), slope main effect (D) and slope variance terms (E-G). Individual plots present the results for high (H) and low (L) levels of among individual (I) and among brood (B) variance. Bar plots present the proportion of simulations that returned a significant effect for a term under true positive (term was non-zero in the simulation) and false positive (term was zero in the simulation) scenarios. In the estimation plot the point is the median and the error bars span the 95% quantile across simulations. The orange dashed line corresponds to the simulated non-zero value, and the green dashed line is at 0 for simulations that correspond to the null hypothesis.

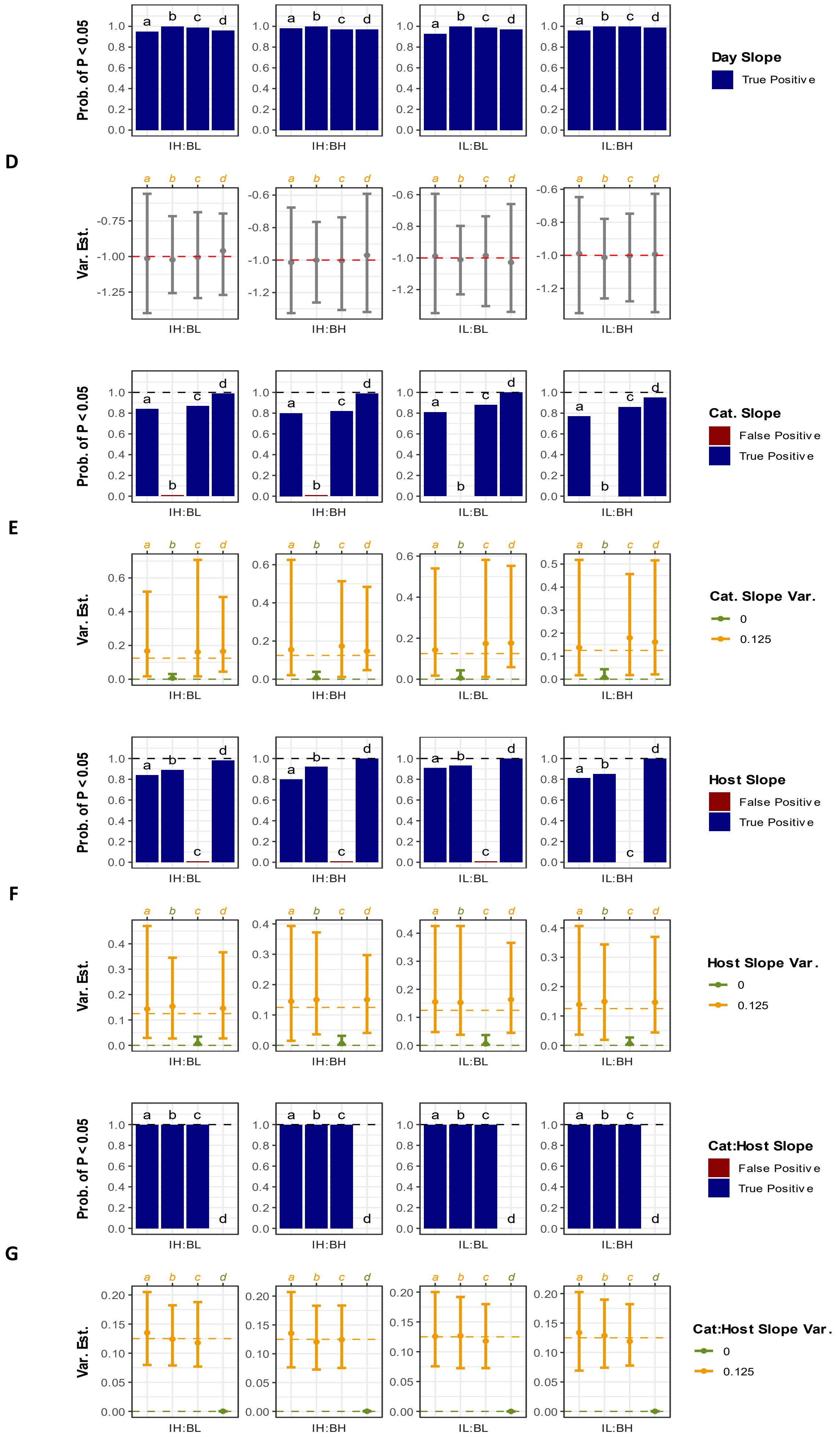
D
E
F
G
Prob. of P < 0.05
Var. Est.
IH:BL
IH:BH
IL:BL
IL:BH
Day Slope
True Positive
Cat. Slope
False Positive
True Positive
Cat. Slope Var.
0
0.125
Host Slope
False Positive
True Positive
Host Slope Var.
0
0.125
Cat:Host Slope
False Positive
True Positive
Cat:Host Slope Var.
0
0.125

**Fig A1 | *cont.***
